# VE-PTP participates in vasculogenic mimicry by preventing autophagic degradation of VE-cadherin

**DOI:** 10.1101/634584

**Authors:** Daniel Delgado-Bellido, Concepción Bueno-Galera, Angel Garcia-Diaz, F. Javier Oliver

## Abstract

Aberrant extra-vascular expression of VE-cadherin has been observed in metastasis associated with Vasculogenic Mimicry (VM); we have recently shown that in VM prone (VM+) tumor cells VE-cadherin is mainly in the form of pVE-cadherin in Y658 allowing an increased plasticity that potentiates VM development. As excessive VE-cadherin phosphorylation is regulated by the phosphatase VEPTP in endothelial cells in the current study we analysed its role in this aberrant phenotype in malignant tumor cells. We show that human malignant melanoma cells VM+, also express VE-PTP although at lower levels than endothelial cells. The complex VE-PTP/VE-Cadherin/p120-catenin act as a safeguard to prevent VE-cadherin degradation by autophagy. Indeed, silencing of VE-PTP results in complete degradation of VE-cadherin with the features of autophagy and increases the global p120 tyrosine phosphorylation status. In summary, we show that VE-PTP is involved in VM formation and disruption of VE-PTP/VE-Cadherin/p120 complex results in enhanced autophagy in aggressive VM^+^ cells.

## Introduction

The term vasculogenic mimicry (VM) describes the formation of perfusion pathways in tumors by highly invasive, genetically deregulated tumor cells: vasculogenic because they distribute plasma and may contain red blood cells and mimicry because the pathways are not blood vessels and merely mimic vascular function. While VM formation is clearly a marker of highly invasive tumor phenotype, mechanisms by which these structures may contribute to adverse outcome are not well understood. It has been proposed that VM formation may facilitate tumor perfusion and the physical connection between VM and blood vessels may also facilitate hematogeneous dissemination of tumor cells. There is a strong association between the histological detection of VM patterns in primary uveal and cutaneous melanomas and subsequent death from metastasis (McLean et al., 1997; Warso et al., 2001), consistent with the in vitro observations that these patterns are generated exclusively by highly invasive tumor cells (Maniotis et al., 1999). ECs express various members of the cadherin superfamily, in particular vascular endothelial (VE-) cadherin (VEC), which is the main adhesion receptor of endothelial adherent junctions. Aberrant extra-vascular expression of VE-cadherin has been observed in certain cancer types associated with VM (Hendrix et al., 2001). VE-PTP (vascular endothelial protein tyrosine phosphatase) is an endothelial receptor-type phosphatase whose name was coined for its prevalence to the bind to VE-cadherin (Nawroth et al., 2002). VE-PTP poise endothelial barrier through helping homotypic VE-cadherin to keep at minimum basal endothelial permeability (Wessel et al., 2014). Knockdown of VE-PTP increases endothelial permeability and leukocyte extravasation. VE-PTP also counterbalances the effects of permeability-increasing mediators such as VEGF, which increase endothelial permeability and leukocyte trafficking, by dephosphorylating VE-cadherin at Tyr658 and Tyr685, leading to stabilization of VE-cadherin junctions (Orsenigo et al., 2012; Wallez et al., 2007).

While studies have focused on the connection between VE-PTP and VE-cadherin in ECs no reports have focused about the role of VE-PTP in VM. We have recently shown that phospho-VEcadherin is highly expressed in VM+ cells and facilitates their pseudoendothelial behaviour by favouring p120/kaiso-dependent gene regulation (Delgado-Bellido et al., 2018). In the current study, we have elucidated a mechanism linking VE-PTP expression to induction of VM in metastatic melanoma through its association with VE-Cadherin/p120 consequent implications in the stability of the complex that promote autophagy. These results place VE-PTP as a dynamic component of VM transformation of melanoma cells owing to its ability retain/safeguard VE-cadherin from being degraded by autophagy in aggressive cells.

## Results and discussion

### VE-PTP expression is essential for VE-cadherin stability and to form VM

Aberrant extra-vascular expression of VE-cadherin has been observed in certain cancer types associated with VM and we have previously shown that most of theVE-cadherin present in VM+ melanoma cells is phosphorylated form in Y658 (Delgado-Bellido et al., 2018). In the current study, we focused in the role of the phosphatase VE-PTP and its interaction with VE-cadherin and its consequences in VM development. Total VE-cadherin and VE-PTP expression was measured in different melanoma cell lines from either cutaneous (C8161, C81-61) or uveal (MUM 2B, MUM 2C) origin as shown in Fig.1A (protein) and b (mRNA). Recently, our group reported that human malignant melanoma cells have a constitutively high expression of pVE-cadherin at position Y658, pV-EC-cadherin Y658 is a target of focal adhesion kinase (FAK) and forms a complex with p120-catenin and the transcriptional repressor Kaiso in the nucleus (Delgado-Bellido et al., 2018). We also have shown that FAK inhibition enabled Kaiso to suppress the expression of its target genes and enhanced Kaiso recruitment to KBS-containing promoters (CCND1 and WNT 11). We observed that silencing of VE-PTP induced a significant reduction of CCND1 and WNT 11 (Kaiso-dependent genes) (Figure 1C) and disrupted VM formation (Fig 1D) suggesting that VE-PTP was also involved in the intracellular dynamic of VE-cadherin resulting in the regulation of kaiso-dependent genes. To evaluate the correlation between the levels of VE-PTP and VE-cadherin we performed a western after siVE-PTP in MUM 2B (Fig 1E) and found an almost complete disappearance of VE-cadherin and phoshoVE-cadherin suggesting that VE-PTP was involved in VE-cadherin stability and needed for phosphoVE-Cadherin to reach the nucleus, as indirectly suggested the results obtained in Fig 1C and D. Interestingly, a completely different situation was found in primary endothelial cells HUVEC where siVE-PTP lead to an accumulation of phosphoVEC in both cytosolic and nuclear compartments (Figure 1F). These results suggested that VE-PTP in malignant melanoma cells was protecting phosphoVEC from degradation.

**Figure 1:**
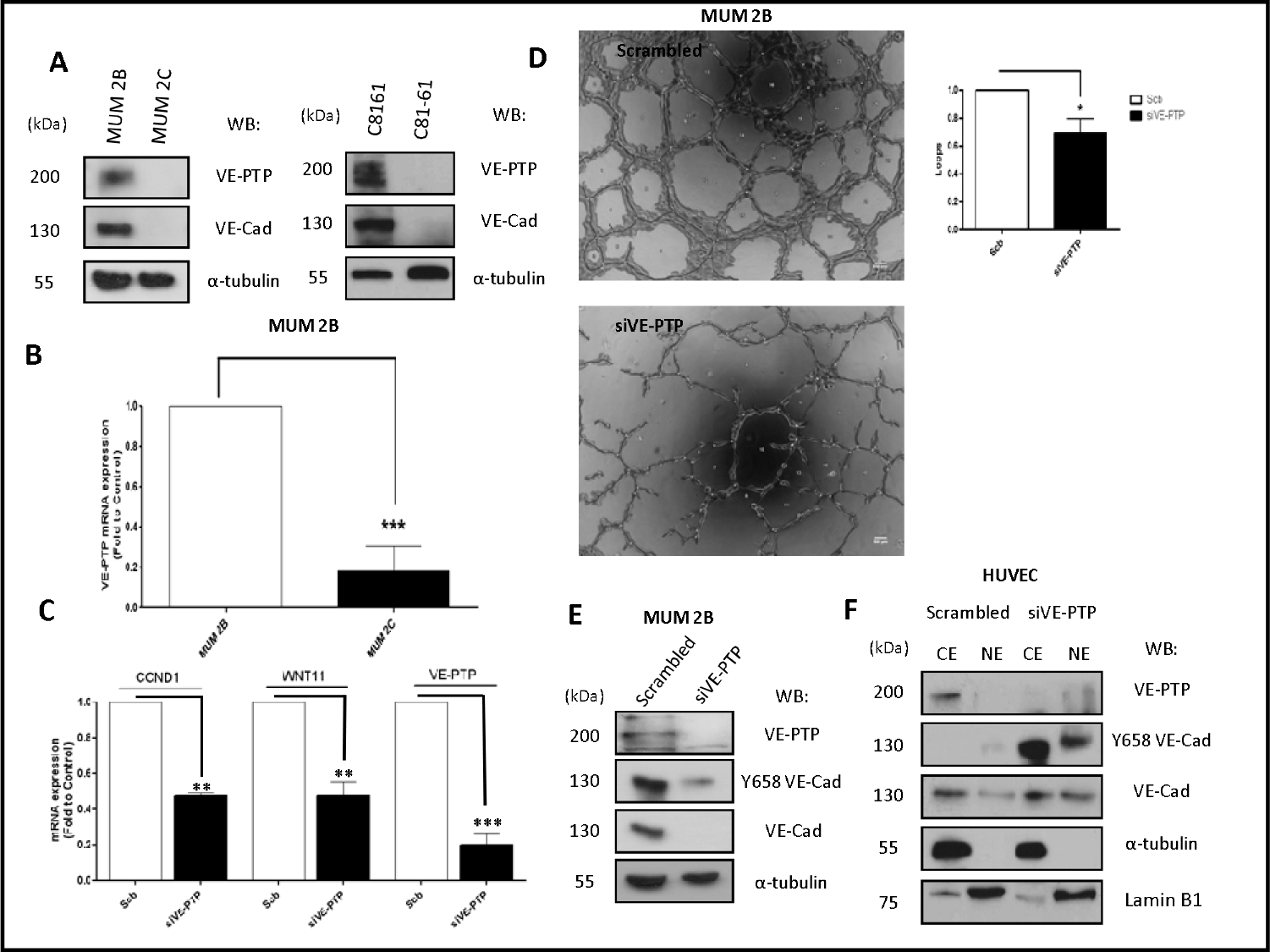
VE-PTP expression is essential for VE-cadherin stability and to form VM: a) The protein expression of VE-PTP is increase in aggressive melanoma cells (MUM 2B and C8161) in compare with poorly melanoma cells (MUM 2C and C81-61), b)RNA expression of VE-PTP also decrease in MUM 2C, c)to see the levels of Kaiso dependent genes that are implicated in VM, qPCR experiments shown that silencing of VE-PTP decrease CCND1 and WNT 11 (Kaiso dependent genes) suggest the possible implications of VE-PTP in VM model. d,e) siVE-PTP abolish the capacity to form VM in aggressive melanoma cells on matrigel, images were acquired using an Olympus CKX41 microscope (bars 500◻μm, the formation of tube-like structures was then quantified by Wimasis program. Each treatment was performed in triplicate, and the experiment was independently repeated at least three times. Results are represented as fold enrichment over input. Asterisks denote significance in an unpaired t test (◻p◻<0.05,◻◻p◻<◻0.01,◻◻◻p◻<◻0.001), and error bars denote SD, f) to confirm the implications of VE-PTP in VM, we performed a siVE-PTP and we show almost total decrease of VE-cadherin and Y658 expression in MUM 2B, g) siVE-PTP on HUVEC in cytosol-nucleus subfracionation experiments increased the phosphorylation of Y658 VE-Cadherin and the consequent internalization into the nucleus,

### VE-Cadherin and VE-PTP form a complex with p120 catenin in melanoma cells

The VE-cadherin-catenin complex provides the backbone of adherens junction in the endothelium. Nonetheless, in non-endothelial cells the proteins interacting with VE-cadherin have not been identified. Using a coIP approach to analyze the VE-cadherin and VE-PTP interacting proteins we shown that VE-Cadherin form a weak complex with VE-PTP in MUM2B (Figure 2A) compared with HUVECs (Fig 2B). Surprisingly, the presence of p120-catenin in complex with VE-PTP was much higher in MUM2B cells as compared to HUVEC cells (Figures 2B and 2C) suggesting that VE-PTP might be involved in the control of p120-catenin phosphorylation status in melanoma cells. To analyze the possible impact of p120 on VE-cadherin stability in MUM2B, we performed a cytosol-nucleus subfractionation assay after silencing p120 and we found that p120 protect the stability of VE-cadherin (Fig 2D). Finally, we performed a coIP of p120 after siVE-PTP in MUM 2B cells, and we observed that binding of VE-cadherin to VE-PTP was lost and resulted in increased global tyrosine phosphorylation of p120, suggesting that VE-PTP safeguard of VE-Cadherin/p120 binding in VM^+^ cells (Fig 2F) and p120-catenin is likely to be a substrate for VE-PTP. These results are compatible with the increased phosphop120 (as result of VE-PTP inactivation) being responsible of complex dissociation to initiate ve-cadherin proteolysis.

**Figure 2:**
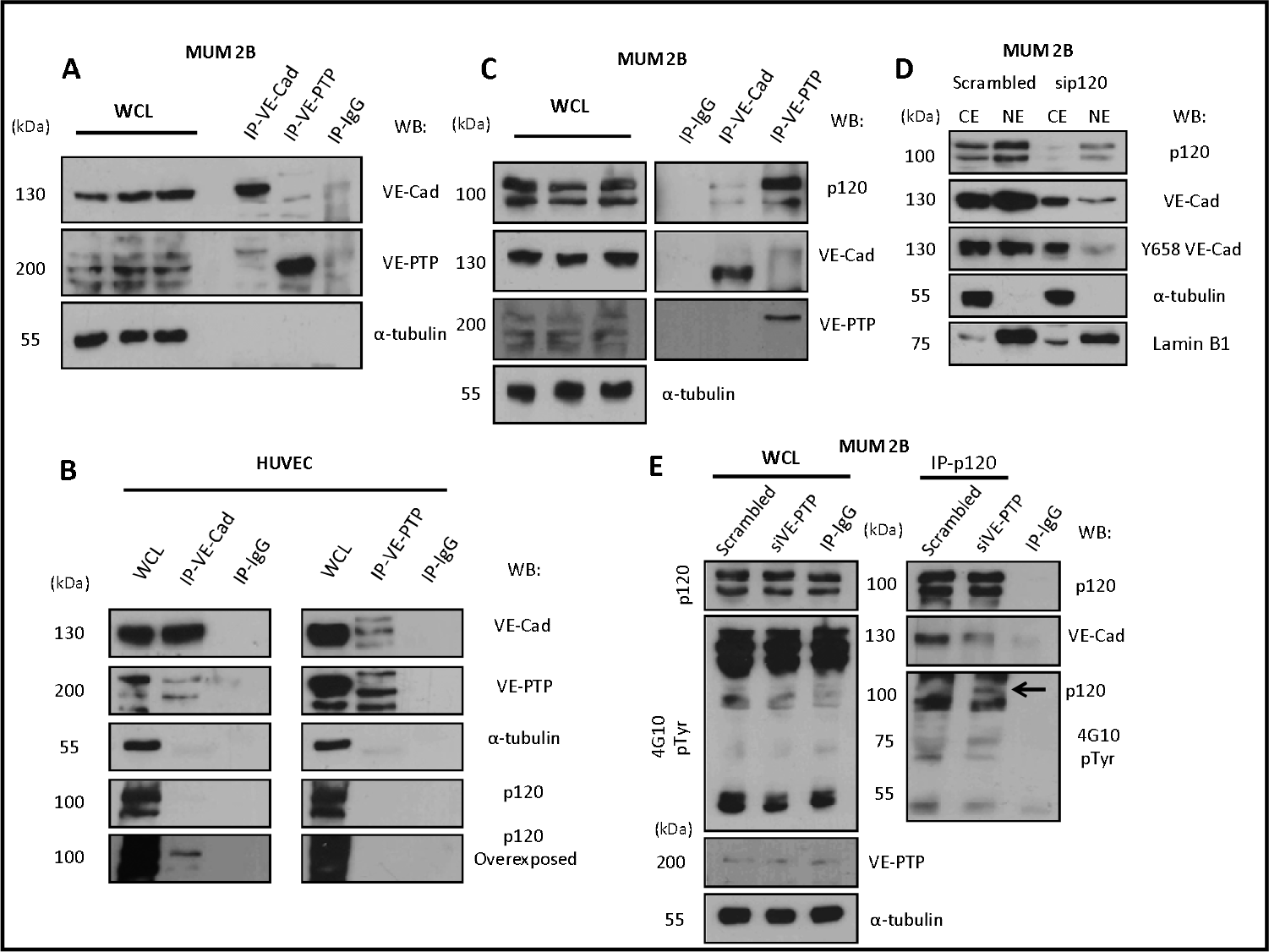
VE-Cadherin and VE-PTP form a complex with p120 catenin in melanoma cells: a) immunoprecipitation of VE-Cadherin and VE-PTP in MUM 2B showed a little union of both proteins in VM cells, b) in contrast, in normal endothelial cells (HUVEC) showed a high union of VE-Cadherin to VE-PTP and only p120 have union with VE-cadherin and not with VE-PTP, c) we want to know what is the union of p120 with VE-cadherin as well as VE-PTP. Surprisingly, p120 have a high union with VE-PTP and VE-cadherin, d) cytosol-nucleus subfracionation with scb and sip120 shown that the stability of VE-cadherin as well the phosphorylation of Y658 is p120-dependent, e) finally, immunoprecipitation of p120 revealed after inhibition of VE-PTP produce a high increase onto p120 phosphorylation and decrease in the union of VE-cadherin.

### VE-Cad/VE-PTP complex dissociation enhanced autophagy

In view of elevated expression of VE-PTP in aggressive melanoma cells and his association with VE-Cadherin/p120 association, we investigated the possible implication of autophagy in these models. To dissect the mechanism leading to VE-Cadherin degradation after the inhibition of VE-PTP, we treat MUM 2B with proteasome inhibitor lactacystin and did not prevent the degradation of VE-Cadherin after VE-PTP disabling (Figure 3A). Macroautophagy (referred to simple as “autophagy”) is a homeostatic “self-eating” pathway that has been conserved among eukaryotic cells. Is a lysosomal-associated process which intracellular components, small portions of cytosol or receives chaperone-associated cargoes are engulfed in double-membrane vesicles, called autophagosomes, to be degraded with lysosomal hydrolyses (Dikic and Elazar, 2018). Recently, different studies report that the formation of VM was promoted by bevacizumab-induced autophagy in GSCs, which was associated with tumor resistance to antiangiogenic therapy through high expression of VEGFR-2 (9). In our setting, inhibition of the fusion of autophagosomes and lysosomes with chloroquine suppressed the degradation of VE-cadherin after siVE-PTP (Fig 3A). Even more, the levels of the mTOR substrate p-p70 (as a readout od mTOR activity and autophagy status) decreased in the MUM 2B K.O cells (Figure 3B), suggesting that VE-cadherin/VE-PTP complex might be restraining autophagy. We then performed an electron microscopy experiment in MUM 2B and C8161 in siVE-PTP or MUM 2B knockout for VE-cadherin; we shown an enhanced autophagic morphology after the VE-PTP silencing or VE-cadherin knockout cells, suggesting that the presence of either protein out of the complex triggered autophagy. To confirm the implication of autophagy on VE-cadherin degradation, we perfomed an experiment to quantify autophagosomes formation under the same conditions described above after transfection LC3-GFP, and we observed that autophagosomes (LC3-GFP puncta) increased following silencing of VE-PTP in both MUM 2B and C8161 VM^+^ cells (Fig 3D). By analysing the cBioPortal database, a platform of 48333 tumour samples, we found that high mRNA levels of PTPRB were inversely associated with the expression of two key genes involved in autophagy, LAMP1 and ATG7 in uveal melanoma (figure S1B), suggesting that in patients this interaction might be relevant to determine autophagy features of the tumor.

**Figure 3:**
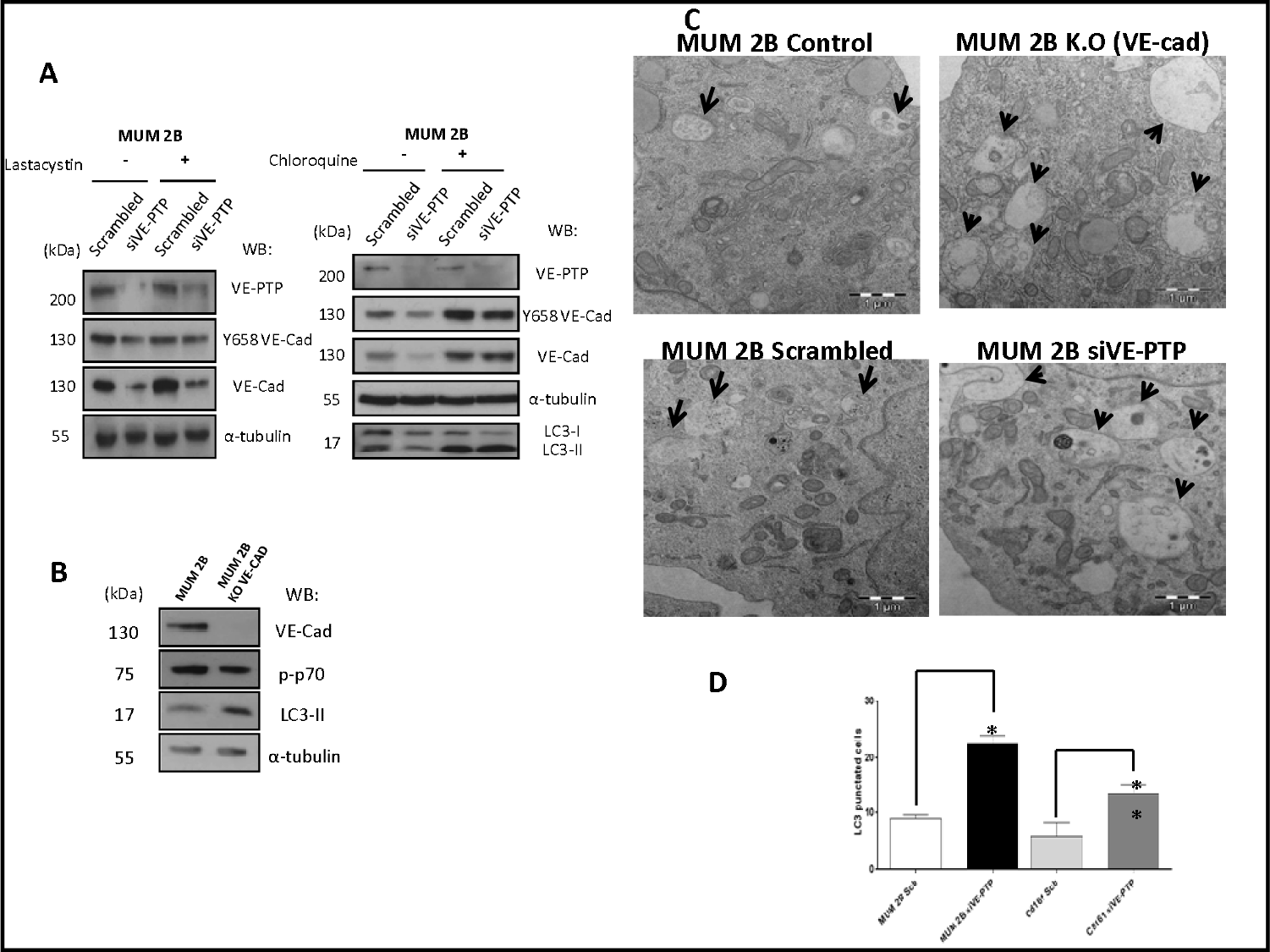
VE-Cad/VE-PTP complex dissociation enhanced autophagy: a) inhibition of proteasome thought lactacystin (30 μM during 30 minutes) with or without siVE-PTP conditions not prevent the VE-cadherin degradation, chloroquine treatment (20 μM during 3h) with the same conditions describe above prevent the degradation of VE-cadherin suggest the possible autophagy implications after siVE-PTP conditions, c) electron microscopy experiments in scrambled, siVE-PTP and MUM 2B K.O conditions with a positive control (HANKs 30 minutes) showed that after inhibition of VE-PTP increase the autophagosomes (bars: 1μm), c) Western blot in MUM 2B K.O cells showed decrease in the expression of p-p70 with an increase in LC3-II formation, d) quantification of autophagosomes (LC3-GFP punctuated cells) increased following silencing of VE-PTP in both MUM 2B and C8161 VM+ cells, each treatment was performed in triplicate, and the experiment was independently repeated at least three times.

While the role of VE-cadherin as a determinant of the pseudo-endothelial behavior of malignant melanoma cells have been widely described, no studies have addressed so far the implications of VE-PTP in VM development. Our previous results have reported that VE-cadherin in VM-prone tumor cells is largely as phosphoVE-cadherin (Y658) and in several intracellular locations (including the nucleus) conferring the cells with the necessary plasticity to undergo pseudo-endothelial differentiation (Delgado-Bellido et al., 2018). To get further information on the cause of these phosphorylated VE-cadherin population we focalized in the phosphatase VE-PTP that keeps VE-cadherin unphosphorylated in endothelial cells. In spite of the large amounts of VE-cadherin, VE-PTP levels in melanoma cells were strongly diminished, suggesting that a majority of the VE-cadherin population is not in complex with VE-PTP (as compared with endothelial cells figures 1F and 2B), then tolerating the accumulation of phospho-VE-cadherin. While endothelial cells tight junctions require a stable and abundant VE-PTP/VE-cadherin complex to keep vascular permeability strictly under control, in melanoma cells the presence of unbound VE-Cadherin to VE-PTP (or the reverse situation) (figure 4) initiates the proteolysis of VE-cadherin trough autophagy. The question still remains how a relatively small amount of VE-PTP protect from proteolysis and what signal emerge from the complex dissociation to activate autophagy. Contrary to endothelial cells, p120-catenin is also strongly attached to VE-PTP (figure 2B and C) and the loss of this complex (after siVE-PTP silencing) leads to p120-catenin increased phosphorylation and VE-cadherin degradation. Globally these results shed light to a new mechanism to control VM through the balance between VE-PTP/VE-cadherin.

**Figure 4:**
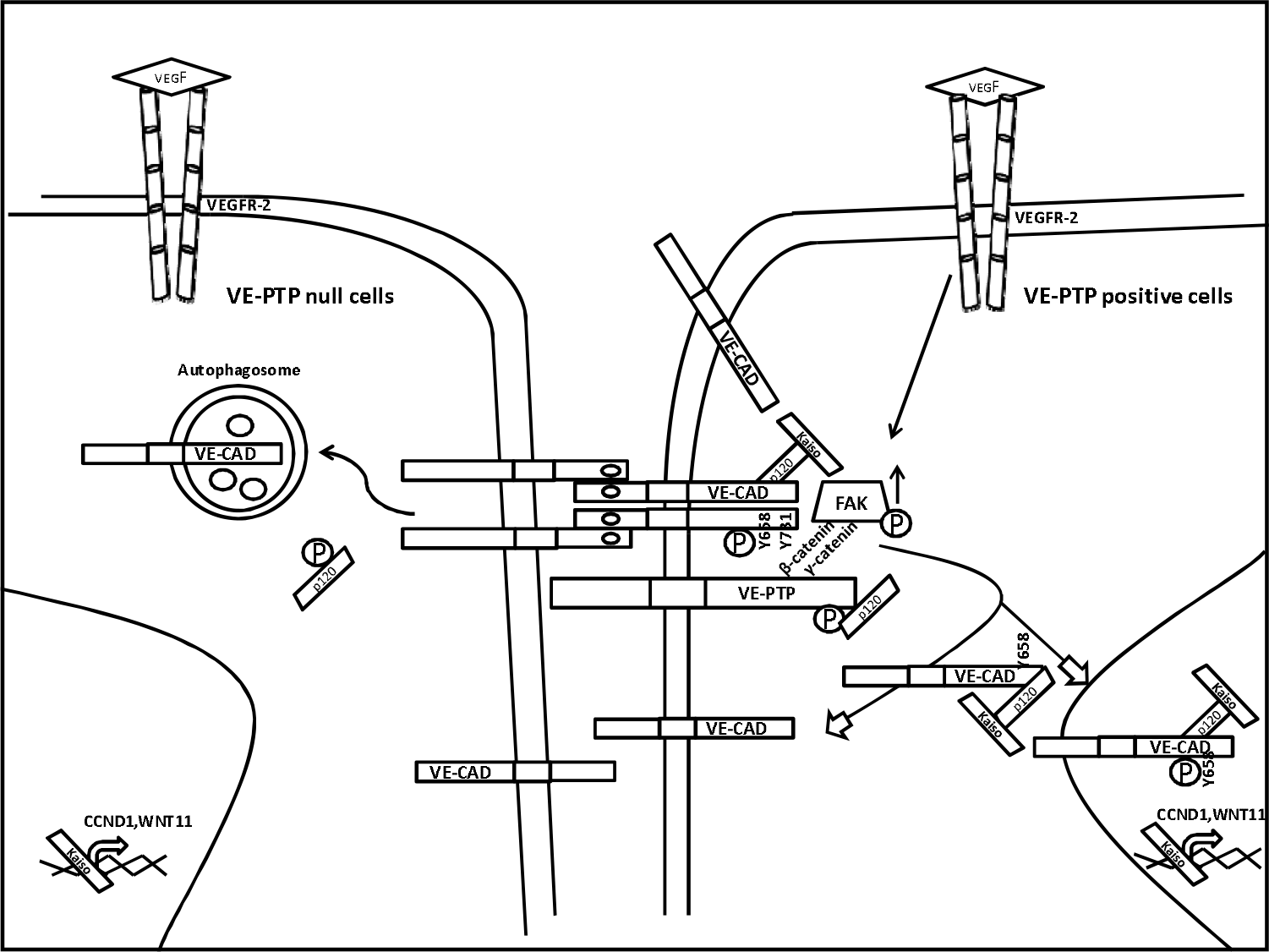
Main signaling pathways involved in vasculogenic mimicry in VE-PTP/VE-Cad null or positive aggressive melanoma cells.

## Materials and methods

### Reagents and antibodies

The following reagents were used: Chloroquine 20 μM during 3h, Lasctacystin 30 μM during 30 minutes. Corning Matrigel Basement Membrane Matrix for in vitro angiogenesis experiments. Antibodies used were: Y658 VEC rabbit (1:1000 WB, 1:100 IF, Thermofisher), VEC C-ter mouse (1:500 WB, 1:50 IF, 2◻μg IP, clone F-8, sc-9989), anti-phosphotyrosine p-Tyr mouse (1:1000 WB, clone 4G10, Millipore), α-tubulin mouse (1:10000 WB, clone B-5-1-2, Sigma-Aldrich), p120 catenin mouse (1:1000 WB, 1:100 IF, 2◻μg IP, BD Biosciences), lamin B1 rabbit (1:1000 WB, Abcam) and VE-PTP mouse (1:1000 WB, 2◻μg IP, Clone 12/RPTPb, BD Biosciences).

### Cell lines

Human melanoma cells MUM 2B, MUM 2C, C8161, and C81-61 were grown in RPMI medium supplemented with 10% fetal bovine serum, 2◻mM of L-glutamine, and 1% penicillin/streptomycin (PAA laboratories). Human umbilical vein endothelial cells (HUVEC) were grown in endothelial cells growth medium-2 (EGM-2) (Lonza). All cells were cultured at 37◻°C and 5% CO2 in incubator cells.

### In vitro angiogenesis assay

The effect of siVE-PTP on the formation of tube-like structures in Matrigel (BD Biosciences) was determined according to manufacturer’s instructions. Briefly, 96-well plates were coated with 50◻μl of BD Matrigel™ Basement Membrane Matrix and allowed to solidify at 37◻°C in 5% CO2 for 30◻min. Cells were treated Scb, siVE-PTP transfected for 48h as described previously. After 48◻h, respectively, of incubation, images were acquired using an Olympus CKX41 microscope. The formation of tube-like structures was then quantified by Wimasis program. Each treatment was performed in triplicate, and the experiments were independently repeated at least three times.

### Quantitative RT-PCR

Total RNA was isolated by RNeasy Mini Kit (Qiagen) according to the manufacturer’s recommendations. About 1◻μg of RNA from each sample was treated with DNase I, RNasinRibonuclease inhibitors (Invitrogen) and reverse-transcribed using iScriptcDNA synthesis kit (Biorad) following the manufacturer’s protocols. cDNA was amplified using the iTaq Universal SYBR green supermix (Biorad). Each reaction was performed in triplicate using CFX96 Real-time PCR detection systems. Primer sequences for the targets and the annealing temperature (60◻°C): 36B4: Forward 5'-CAGATTGGCTACCCAACTGTT-3', Reverse 5'-GGCCAGGACTCGTTTGTACC-3, CCND1: Forward 5'-CCGTCCATGCGGAAGATC-3' Reverse 5'-GAAGACCTCCTCCTCGCACT-3'; WNT11: Forward 5'-GCTTGTGCTTTGCCTTCAC-3', Reverse 5'-TGGCCCTGAAAGGTCAAGTCTGTA-3', VE-PTP: Forward 5'-TGCTAAGTGGAAAATGGAGGCT-3', Reverse 5'-GCCCACGACCACTTTCTCAT-3'.

### Gene editing

MUM2B knockout (ko) cells for the VE-Cad gene were generated using the CRISPR-Cas9 technology. Five different sgRNAs were designed using the Zhang Lab Optimized CRISPR design tool and cloned into the pL-CRISPR.EFS.GFP which was purchased from the Addgene public repository (#57818). sgRNA guides were validated in HEK293T Cells using the GeneArt Genomic Cleavage Detection Kit (Invitrogen, Carlsbad, USA) according to the manufacturer’s instructions. Lentiviral particles for the best two sgRNAs in terms of allelic disruption (GGCAGGCGCCCGATGTGGCG and GATGATGCTCCTCGCCACATC).

### Transfection of small interfering siRNA

Cultured cells were transiently transfected with an irrelevant siRNA (5'-CCUACAUCCCGAUCGAUGAUG-3') 50nM. siVE-PTP: 5'-GACAGUAUGAGGUGGAAGU-3', 50nM, sip120: 5'-GGATCACAGTCACCTTCTA-3', 50nM, were transfected for 48 hours using JetPrime (Polyplus transfection) according to the recommendations.

### Immunobloting, immunoprecipitation, subfractionation cytosol-nucleus

For simple coimmunoprecipitation, cells were lysed in lysis buffer (50 mM Tris/HCl ph 8, 120 mM NaCl, 0,1 % NP-40, 1mM EDTA, 10 mM NaF, 1 mM Na3VO4 and supplemented with a protease inhibitor cocktail (1 tablets to 10 ml of lysis buffer, Roche) for 30 minutes at 4°C. Lysates were cleared by 13.000 rpm centrifugation for 10 minutes at 4°C and incubated over night at 4°C with respective antibodies. Consequently, the next day IP lysates were incubated for 2 hours at 4°C with 50 μl of of Dynabeads™ Protein G for Immunoprecipitation (ThermoFischer). Dynabeads were washed 3 times with low salt 120 mM lysis buffer and 2 times with high salt 300 mM lysis buffer. All lysates were separated by dodecyl sulfate-polyacrylamide (7,5%, Biorad) gel electrophoresis and transferred to PVDF membrane (Pall laboratory) by semi-wet blotting.

Accordingly with the article of Maxie Rockstroh et al in 2011, for subfractionation cytosol-nucleus, cells were lysed in lysis buffer (250 mM sucrose, 50 mM Tris-HCl ph 7,4, 5 mM MgCl2, 1 mM Na3VO4, 0,25 % NP-40 and supplemented with a protease inhibitor cocktail (1 tablets to 10 ml of lysis buffer, Roche) for 10 minutes at 4°C, lysated were centrifuged at 500 g for 5 minutes, supernatant was considerate cytosolic fraction, pellet was resuspended in buffer 2 (1M sucrose, 50 mM Tris-HCl ph 7,4, 5 mM MgCl2) and centrifuged at 3000 g for 5 minutes, supernatant was discarded. Pellet was resuspended in nuclear buffer (20 mM Tris-HCl ph 7,4, 0,4 M NaCl, 15% glycerol, 1,5% Triton X-100) for 45 minutes at 4°C in agitation. This lysate were centrifuged at 5000 g for 5 minutes, supernatant was considered nuclear fraction.

### Electron microscopy

The MUM 2B extracted were washed with PBS, prefixed for 30 min in a fixation solution (0.1 M cacodilate buffer pH 7.4 and osmium tetraoxyde) for 60 min at 4 °C. After this treatment, tissues were washed with MilliQ water and the samples were stained with uranil acetate. The ultrathin sections were cut with a diamond knife in an ultramicrotome (Reichert Ultracut S). The samples were analyzed in a TEM Zeiss 902 with 80 KV of voltage acceleration (CIC-UGR).

### Autophagy assay

GFP-LC3-expressing cells have been used to demonstrate the induction of autophagy. The GFP-LC3 expression vector was kindly supplied by Dr T Yoshimori (National Institute for Basic Biology, Okazaki, Japan). MUM 2B and C8161 were transiently transfected (0,5 μgr) with this vector together with jetPrime (Polyplus transfection, Illkirch, France) according to the manufacturer's protocol. The assay was performed on cells grown in six-well plates and after the different treatment with HANK buffer 10 minutes. To determine LC3 localization, GFP-LC3-transfected cells were observed under a Zeiss fluorescence microscope. To determine LC3-II translocation, we performed western blot of LC3-I and its proteolytic derivative LC3-II (18 and 16 kDa, respectively) using a monoclonal antibody against LC3 (NanoTools, clone 5F10, Ref 03231-100/LC3-5F10).

## Acknowledgements

This work was supported by the grants from Spanish Ministry of Economy and Competitiveness SAF2015-70520-R, and Spanish Ministry of Science and Technology RTI2018-098968-B-I00 and CIBERONC ISCIII CB16/12/00421 to FJO.

## References

Delgado-Bellido, D., Fernandez-Cortes, M., Rodriguez, M. I., Serrano-Saenz, S., Carracedo, A., Garcia-Diaz, A. and Oliver, F. J. (2018). VE-cadherin promotes vasculogenic mimicry by modulating kaiso-dependent gene expression. Cell Death Differ.

Dikic, I. and Elazar, Z. (2018). Mechanism and medical implications of mammalian autophagy. Nat Rev Mol Cell Biol 19, 349–364.

Hendrix, M. J., Seftor, E. A., Meltzer, P. S., Gardner, L. M., Hess, A. R., Kirschmann, D. A., Schatteman, G. C. and Seftor, R. E. (2001). Expression and functional significance of VE-cadherin in aggressive human melanoma cells: role in vasculogenic mimicry. Proc Natl Acad Sci U S A 98, 8018–23.

Maniotis, A. J., Folberg, R., Hess, A., Seftor, E. A., Gardner, L. M., Pe'er, J., Trent, J. M., Meltzer, P. S. and Hendrix, M. J. (1999). Vascular channel formation by human melanoma cells in vivo and in vitro: vasculogenic mimicry. Am J Pathol 155, 739–52.

McLean, I. W., Keefe, K. S. and Burnier, M. N. (1997). Uveal melanoma. Comparison of the prognostic value of fibrovascular loops, mean of the ten largest nucleoli, cell type, and tumor size. Ophthalmology 104, 777–80.

Nawroth, R., Poell, G., Ranft, A., Kloep, S., Samulowitz, U., Fachinger, G., Golding, M., Shima, D. T., Deutsch, U. and Vestweber, D. (2002). VE-PTP and VE-cadherin ectodomains interact to facilitate regulation of phosphorylation and cell contacts. EMBO J 21, 4885–95.

Orsenigo, F., Giampietro, C., Ferrari, A., Corada, M., Galaup, A., Sigismund, S., Ristagno, G., Maddaluno, L., Koh, G. Y. and Franco, D. (2012). Phosphorylation of VE-cadherin is modulated by haemodynamic forces and contributes to the regulation of vascular permeability in vivo. Nature communications 3, 1208.

Wallez, Y., Cand, F., Cruzalegui, F., Wernstedt, C., Souchelnytskyi, S., Vilgrain, I. and Huber, P. (2007). Src kinase phosphorylates vascular endothelial-cadherin in response to vascular endothelial growth factor: identification of tyrosine 685 as the unique target site. Oncogene 26, 1067–77.

Warso, M. A., Maniotis, A. J., Chen, X., Majumdar, D., Patel, M. K., Shilkaitis, A., Gupta, T. K. and Folberg, R. (2001). Prognostic significance of periodic acid-Schiff-positive patterns in primary cutaneous melanoma. Clin Cancer Res 7, 473–7.

Wessel, F., Winderlich, M., Holm, M., Frye, M., Rivera-Galdos, R., Vockel, M., Linnepe, R., Ipe, U., Stadtmann, A., Zarbock, A. et al. (2014). Leukocyte extravasation and vascular permeability are each controlled in vivo by different tyrosine residues of VE-cadherin. Nat Immunol 15, 223–30.

